# Therapeutic relevance of NLPA Lipoprotein to combat biofilm-associated infections in Acinetobacter baumannii

**DOI:** 10.64898/2026.05.18.725845

**Authors:** Umarani Brahma, Sharon Munagalasetty, Vasundhra Bhandari

## Abstract

*Acinetobacter baumannii* is a leading multidrug-resistant critical priority pathogen in healthcare settings, where biofilm formation confers survival and antibiotic tolerance. Targeting virulence associated proteins offers an alternative to conventional bactericidal strategies. Here, the inner membrane anchored lipoprotein NLPA, implicated in biofilm associated adaptation, was studied as a putative anti-virulence target using an integrated in silico pipeline and complementing the computational findings. The Alpha fold-derived structure of NLPA served as the basis for virtual screening of approximately 1.6 million compounds, with subsequent prioritization guided by MM/GBSA calculated binding free energies to highlight the top promising candidates. Molecular dynamics simulations demonstrated stable NLPA ligand complexes, as indicated by equilibrated RMSD, low residue fluctuations in the binding region, and persistent interaction networks over time. Pharmacokinetic evaluation indicated that the compounds satisfied Lipinski’s Rule of Five and had overall acceptable ADMET characteristics. Two compounds, NLPA-6 and NLPA-3, showed the most favourable predicted binding free energies, suggesting strong and stable interactions within the NLPA binding site. NLPA-3 was evaluated in vitro against *A. baumannii* to validate the computational outcomes. The compound displayed moderate antibacterial activity with a MIC of 125 μg/mL and demonstrated 55.75% inhibition of biofilm formation at 4x MIC. In addition, in macrophage infection studies, NLPA-3 decreased intracellular bacterial survival to 19.25% at 50 μg/mL, suggesting that it may disrupt virulence pathways linked to persistence. In whole, these findings identify promising NLPA targeting compounds and support the feasibility of NLPA as an anti-virulence target.

## 1. Introduction

Globally, multi-drug-resistant gram-negative bacterial infections surge in mortality and morbidity rates, presenting a serious threat to human health. Their exceptional ability to rapidly acquire and disseminate antimicrobial resistance (AMR) determinants amplifies their clinical impact on modern healthcare (Macesic, Uhlemann et al. 2025). These bacteria pose a profound and escalating threat to public health and ecological balance, demanding immediate, sustained, and globally coordinated action (Subhasmita Mallik 2025). One of the biggest hazards related to AMR worldwide is *Acinetobacter baumannii*, which has been declared as a critical priority pathogen by the World Health Organization (WHO) in the revised 2024 Bacterial Priority Pathogens List (BPPL) (WHO 2024). This bacterium is closely associated with life-threatening hospital-acquired infections, including bacteremia, pneumonia, and urinary tract or skin and soft tissue infections (Maure, Robino et al. 2023). Globally, carbapenem-resistant *A. baumannii* infections have risen to 30%, with high prevalence, 80% seen in many regions of Asia and 50% in southern and eastern Europe (Macesic, Uhlemann et al. 2025). The rising incidence of infections due to carbapenem, colistin and the novel siderophore, cefiderocol resistance presents consequential treatment challenges (Scribano, Cheri et al. 2024). The antibiotic resistance and biofilm-forming abilities of the bacteria make it a therapeutic challenge, especially in healthcare settings where patients are at risk of developing long-term infections (Upmanyu, Haq et al. 2022).

*A. baumannii* stands out exceptionally for its extraordinary capacity to acquire a plenitude of resistance factors and uses its virulence factor suite to withstand a various host and environmental stresses, which makes the eradication of infection a formidable challenge. Among these virulence determinants, outer membrane proteins (OMPs) represent a prominent and functionally versatile group that contribute significantly to bacterial fitness and pathogenicity (Uppalapati, Sett et al. 2020).

These OMPs are one of the prominent categories of virulence determinants that have attracted a lot of scientific interest due to their unique location, extensive distribution, and vital functions in bacterial physiology and pathogenicity (Uppalapati, Sett et al. 2020). In Gram-negative bacteria outer membrane structural and functional integrity is maintained in large part by OMPs (Zhao, Hu et al. 2024). Due to their crucial role bacterial pathogenesis these represent potential prospects for therapeutic intervention and vaccine development because of their varied roles in disease persistence (Uppalapati, Sett et al. 2020).

Among all these virulence factors, OmpA has received the greatest amount of attention due to its central function in controlling *A. baumannii* adhesion, aggression, biofilm formation, outer membrane vesicle (OMV) production, inactivation of the complement cascade and host immune response and death (Nie, Hu et al. 2020) (Kim, Kim et al. 2019). However, there remain other less characterized membrane associated proteins with critical function and therapeutic relevance.

Among these, NLPA (new lipoprotein A) represents one such protein which was first identified in *E. coli* where its deletion showed no remarkable difference in cell growth, morphology or chemical sensitivity, considering it as non-essential. It is a periplasmic protein or inner membrane-anchored lipoprotein, involved in the import of methionine (Bodero, Pilonieta et al. 2007). Another study identified it as a key antigenic factor involved in the biogenesis of outer membrane vesicles. Few studies have reported that the production of outer membrane vesicles is reduced when *NLPA* is disrupted. A critical transcriptional regulator in biofilm formation, CsgD regulates NLPA in *E. coli*. Expression of CsgD starts in the mid-logarithmic phase and continues until the stationary phase, thereby ensuring sustained regulation of genes for persistence (Schwechheimer and Kuehn 2013). Studies on the *A. baumannii* reported that recombinant expression of NLPA effectively elicited humoral and cellular immunological responses in BALB/c mice. It has been implied from the previous study that NLPA might be a DNA vaccine candidate against *A. baumannii*, since elevated levels of IgG, IgM, IFN-γ, IL-2, IL-4, and IL-12 indicated significant immunogenicity (Hashemzehi, Doosti et al. 2018). In *A. baumannii*, several outer membrane and periplasmic proteins have been explored as subunit vaccine candidates, reflecting their roles in pathogenesis and immune recognition (Jeffreys, Chambers et al. 2022). Expression of NLPA was also found to be upregulated in strong biofilm-forming A. *baumannii* clinical isolates (Umarani Brahma 2025) (*bioRxiv* (preprint)). Given its role in OMV biogenesis and biofilm-associated pathogenicity, NLPA emerges as a multifaceted determinant of *A. baumannii* pathogenicity. Its dual role in bacterial pathogenicity not only represents a promising vaccine candidate but also a potential drug target. To reduce bacterial pathogenicity, virulence-associated proteins may potentially be investigated as anti-virulence targets. Anti-virulence approaches interfere with mechanisms necessary for infection, including adhesion, invasion, and stress adaptability, rather than inhibiting bacterial growth. In order to combat multidrug-resistant bacteria, these methods provide potential alternative or supplementary therapy by disarming pathogenic systems without applying substantial bactericidal pressure (Li, Jia et al. 2025).

A growing number of researchers are turning to computer-aided drug discovery (CADD) methods because they overcome the time and monetary limits as faced by traditional experimental approaches. These include target identification, virtual screening of huge libraries for lead compounds, optimization of promising hits, and in silico assessment of their toxicity (Wankhede 2024). These also provide a quick and economical way of predicting and evaluating probable drug-target associations before experimental confirmation, which expedites the discovery pipeline (Antunes 2021). In the present study, a comprehensive *in-silico* screening of diverse compound libraries to identify inhibitors targeting NLPA as a potential anti-virulence therapeutic strategy against multidrug-resistant *A. baumannii*.

## 2. Materials and Methods

### a. Protein structure optimization and validation

Since the Protein Data Bank (PDB) did not have any specific information about the crystallized structure of the *A. baumannii* NLPA protein, the Alpha fold was used to predict its 3D structure. The protein’s amino acid sequence (Uniprot ID: D0CE28) was retrieved from the UniProt database (Ros-Lucas, Martinez-Peinado et al. 2022). Maestro was used to view the three-dimensional protein structure. Following that, the built model was pre-processed using the protein preparation wizard of Schrodinger Suite 2024, which refined the protein structure for docking by setting bond orders, filling in missing loops, adding hydrogens, and deleting water molecules that are more than 3Å distances from the protein (Sahayarayan, Rajan et al. 2021). Along with that, the protein was optimized by using PROKA at a pH of 7.4 while generating het states using the Epik module. Subsequently, an OPLS4 force field was used to minimize the energy of the protein structure (Peele, Potla Durthi et al. 2020) (Schrödinger Release 2022-4: Protein Preparation Workflow, Schrödinger, LLC, New York, NY, 2022).

### b. Binding site mapping and grid generation

The binding site of the NLPA protein was predicted and evaluated using the Site Map module of the Maestro suite. We used the site score and the D score (druggability score) to assess the binding site. The D score will predict the target’s druggability assessment and the site score to evaluate the ligand’s likelihood of binding to that site. After identifying every possible receptor binding site, it was configured to set a minimum of fifteen site points per site. Using the ‘receptor grid generating panel,’ the grid coordinates around the binding area were specified, thereby defining the ligand binding site. This defines the active site’s size and position and represents the receptor’s shape and properties more accurately. For the NLPA receptor, van der Waals radii were applied during model preparation, employing a partial charge cut-off of 0.25 Å and a default scaling factor of 1.0 Å to refine the binding site representation (Halgren 2009). (Schrödinger Release 2022-4: Site Map, Schrödinger, LLC, New York, NY, 2022).

### c. Ligand acquisition and Virtual screening

A comprehensive dataset of around 1.6 million small molecules was compiled from various chemical libraries, such as VITAS-M, Med Chem Express, and Selleck Chem. These compounds were retrieved in Structural Data Format (SDF) to continue virtual screening analyses. They included a broad range of chemical classes and libraries, from drugs approved by the FDA and Phase I clinical candidates to antibiotics, alkaloids, repurposed molecules, and novel or uncharacterized compounds. The screening of these libraries was carried out using the glide virtual screening workflow (vsw) panel. The ligand screening was carried out systematically, with duplicates being eliminated and ligands being filtered using Lipinski’s rule. Epik was used to produce various ionization states at a pH of 7.0 ± 2.0, leaving all other parameters left at their default values. In order to conduct molecular docking, the generated grid file was included. Three distinct docking modalities were executed on the receptor site created by the grid: high throughput virtual screening (HTVS), standard precision (SP) and extra precision (XP). Only the top-performing candidates from the HTVS phase underwent XP docking for an improved assessment of binding interactions and precise prediction of final poses in order to maximize computational efficiency. The glide energy and XP glide rescoring calculations were applied to generate the primary ligand-receptor complexes. The interactions between the receptor and the ligand were observed using the 2D viewer (Owoloye, Ligali et al. 2022). (Schrödinger Release 2022-4: Glide, Schrödinger, LLC, New York, NY, 2022).

### d. ADME properties and Drug likeliness

Using the Qik Prop module of Maestro, ADME analysis was carried out on the top hits. Many important physicochemical and pharmacokinetic parameters were calculated, including their molecular weight (mol_MW), number of hydrogen bond acceptors (accptHB), number of hydrogen bond donors (donorHB), predicted aqueous solubility (QPlogS), predicted octanol/water partition coefficient (QPlogPo/w), predicted apparent Caco-2 cell permeability (QPPCaco), predicted brain/blood partition coefficient (QPlogBB), percent human oral absorption, and predicted metabolic reactions (#metab). The Lipinski rule of five, which is used for determining the pharmacokinetic profile of drugs, is also assessed (Ntie-Kang, Mbah et al. 2013) (Schrödinger Release 2022-4: QikProp, Schrödinger, LLC, New York, NY, 2022).

The toxicity of the desired compounds was assessed using the ProTox-3.0 web server. Compound IDs, whether by name or SMILES (Simplified Molecular-Input Line-Entry System) strings, were provided for predicting probable toxicities based on their chemical structure. The analysis predicted organ toxicity and primary toxicity endpoints, such as carcinogenicity, immunotoxicity, mutagenicity, and cytotoxicity, along with Tox21 nuclear receptor signaling and stress response pathways. Predictions were produced for both acute toxicity and toxicity endpoints.

The output included the three most structurally analogous drugs having confirmed rat oral toxicity predicted median lethal dosage (LD_50_, mg/kg body weight), toxicity classification, prediction accuracy, average structural similarity, and the three most structurally analogous drugs having experimentally confirmed rat oral toxicity data. Toxicity categorization was conducted in accordance with the Globally Harmonized System (GHS). Compounds having an LD_50_ of 5 mg/kg or less were classified as Class 1, while those with LD_50_ values ranging from 5 to 50 mg/kg were classified as Class 2; both classifications are deemed lethal. Class 3 substances (LD_50_: 50–300 mg/kg) are deemed harmful upon ingestion, but Class 4 and Class 5 compounds (LD_50_: 300–5000 mg/kg) have somewhat reduced toxicity. Compounds classified as Class 6, with LD_50_ values more than 5000 mg/kg, are deemed non-toxic (Banerjee, Kemmler et al. 2024).

### e. Prime MM/GBSA Analysis

The MMGBSA (molecular mechanics generalized Born surface area) calculation yielded the relative binding free energy (ΔG bind) of each ligand molecule to assess ligand binding affinity with the receptor using the Prime module. The calculations of free binding energy for the ten most promising compounds were conducted using the VSGB 2.1 solvation model and the OPLS4 force field. The equation for calculating binding affinity is as follows:

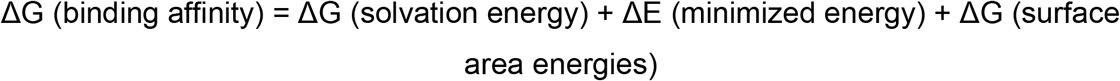

Where, ΔG (solvation energy) represents the difference between the solvation energy of GBSA for the NLPA–inhibitor complex and the total of the solvation energies for unbound NLPA and its associated inhibitor. ΔE (minimized energy) represents a difference between the energy of the NLPA–inhibitor complex and the cumulative energies of the unbound NLPA with its corresponding inhibitor. ΔG (surface area energies) represents the difference between the surface area energy of the NLPA– inhibitor complex and the cumulative energies of unligated NLPA and the inhibitor (Halder, Das et al. 2022). (Schrödinger Release 2022-4: Prime, Schrödinger, LLC, New York, NY, 2022).

### f. Interaction fingerprint analysis of ligand-protein complexes

Structural Interaction Fingerprint (SIFt) analysis was utilized to analyze the common binding pattern of the docking poses to identify residues that make polar or non-polar interactions. SIFt depicts the intermolecular interactions of 3D protein-inhibitor complex by generating a binary interaction fingerprint. SIFt analysis was performed using Interaction Fingerprint Panel (Schrödinger Release 2022–4: Canvas, Schrödinger, LLC, New York, NY, 2022) using docked poses. The fingerprints encode the ligand’s spatial relationship with the protein and the interacting residue types. The information includes the nature of ligand–protein interactions, like hydrogen bonding, hydrophobic contacts, electrostatic forces, and van der Waals interactions, along with identification of key binding-site residues like hydrophobic, aromatic, charge, polar, side chain, and backbone essential for these interactions (Karpurapu, Kakarala et al. 2024).

### g. Molecular dynamics (MD) simulation

Unlike biological processes that include dissolving the protein and ligand in water, molecular docking does not precisely replicate these events. The stability of the hits obtained from virtual screening was assessed using the MD module. Thus, MD simulations were used to analyze the non-bonding interactions between the ligand and the protein over a duration of 100 ns, aimed at understanding the stability of the most promising leads identified. Firstly, a membrane was set to the protein-ligand complex using the system builder module and positioned within the orthorhombic box with a solvent layer that was 10 Å thick using the TIP3P solvent model. The OPLS4 force field was prepared and further neutralized by adding 0.15 M NaCl to the buffer. Additionally, minimization was done using the minimizing tool. Using an NPγT, ensemble class which maintains a constant temperature throughout the simulation together with constant normal pressure and lateral surface tension of membranes (constant pressure temperature) constant pressure ensemble at constant volume, the MD simulation was run at 300 K and 1.01325 bar of pressure for 100 ns. With the trajectory’s recording interval (ps) set at 100 frames per second, around 1000 frames were produced throughout the MD simulation. The simulation interaction diagram (SID) tool included with Desmond was then used to generate reports (Bowers 2006). (Schrödinger Release 2024: Desmond Molecular Dynamics System, D. E. Shaw Research, New York, NY, 2024).

### h. Cross-correlation motion analysis

The dynamic cross-correlation matrix (DCCM) is a 3D matrix representation derived from atomic fluctuations which captures coherent and long-range residue motions, where diagonal elements reflect self-correlation and off-diagonal elements represent inter-residue coupling. DCCM was computed to analyze time-dependent correlated motions of protein residues by using trj_essential_dynamics.py script with - cross_correlation flag on MD trajectories (Qiao, Wang et al. 2024).

### i. Susceptibility and anti-biofilm activity of NLPA-3 in *A*.*baumannii*

In a 96-well plate format, the micro-broth dilution assay was used to determine the minimum inhibitory concentrations (MICs). The micro-broth dilution experiment with concentrations ranging (0 – 512 ug/ml) was carried out in accordance with the Clinical and Laboratory Standards Institute (CLSI) standards with minimal modifications using the resazurin dye.

The inhibitory effect of NLPA-3 on biofilm formation by *A. baumannii* BAA-747 was evaluated using crystal violet (CV) assay. An overnight culture diluted in 1:200 in tryptic soy broth (TSB) supplemented with 2% glucose was dispensed into a 96-well plate (200 µL/well). The plate was incubated at 37 °C for 24 h to allow biofilm formation, followed by three times of gentle washing with PBS to remove planktonic cells. After washing, varying concentrations of NLPA-3 ranging from 0 to 500 µg/mL were added, and the plate was further incubated at 37 °C for 16 h. The wells were subsequently washed three times with PBS and fixed with methanol for 15 min. After air drying for 30 min, 0.1% crystal violet solution was added to each well and incubated at room temperature for an additional 20 min. Excess stain was removed by washing with distilled water, and the bound dye was solubilized using 33% acetic acid. Biofilm biomass was quantified by measuring absorbance at 590 nm. The mean absorbance values of treated wells were compared with untreated controls to determine the percentage inhibition of biofilm formation (Brahma, Kothari et al. 2018).

### j. Assessment of intracellular survival in macrophages and cytotoxicity

RAW 264.7 murine macrophage cells were maintained in DMEM supplemented with 10% fetal bovine serum (FBS) and antibiotics. For intracellular survival assays, cells were seeded in 24-well tissue culture plates at a density of 5 × 10^5^ cells per well in antibiotic-free medium and incubated at 37 °C in a 5% CO_2_ atmosphere. The macrophages were infected with selected strong and weak biofilm-forming *A. baumannii* clinical isolates at a multiplicity of infection (MOI) of 10 (5 × 10^6^ CFU/mL) and incubated for 2 h. Following infection, cells were washed three times with 1× PBS to remove non-adherent bacteria, and gentamicin (200 µg/mL) was added for 1 h to eliminate extracellular bacteria. The infected macrophages were subsequently washed with 1x PBS and treated with meropenem and NLPA-3 at concentrations of 0, 0.2, 10, and 50 and 100 µg/mL for an additional 24 h under the same incubation conditions. After treatment, cells were washed twice with 1x PBS and lysed using 0.1% saponin. The lysates were serially diluted and plated on tryptic soy agar (TSA) plates for enumeration of intracellular bacterial counts. The results were expressed as percentage bacterial survival relative to untreated control cells (Brahma, Kothari et al. 2018).

## 3. Results and Discussion

### a. Target protein preparation and validation

NLPA is an inner-membrane-anchored lipoprotein reported to have a role in methionine import and is found to play a role in biofilm production (Schwechheimer and Kuehn 2013, Hashemzehi, Doosti et al. 2018). The projected 3D model was obtained from AlphaFold Protein Structure, since the Protein Data Bank (PDB) does not include an experimentally verified three-dimensional structure of NLPA. The structure of NLPA, which has 292 amino acid residues, was then refined using energy minimization (Figure 1A). Structural validation using a Ramachandran map (Figure 1B).

**Figure 1.**
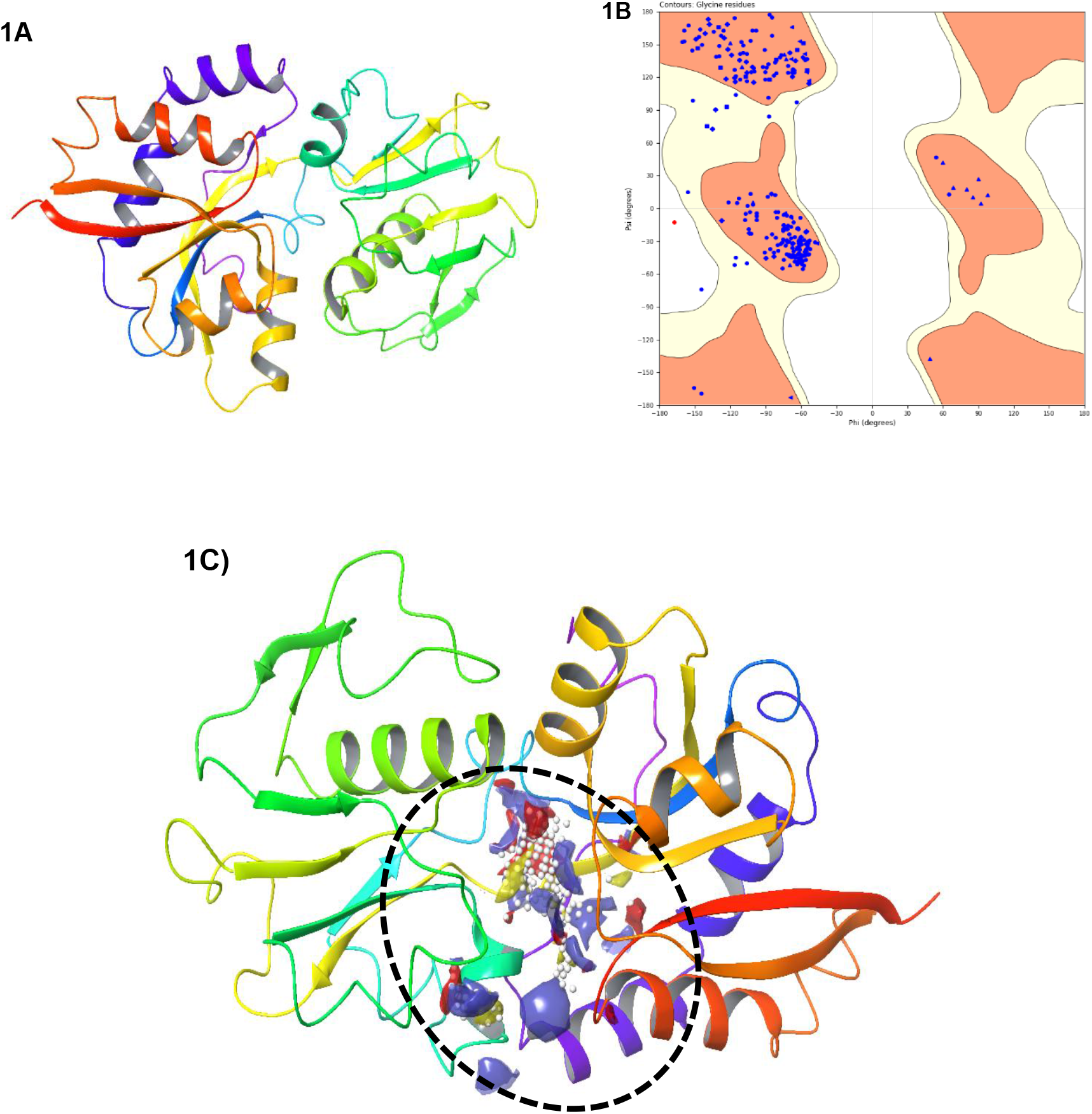
A-C: Structural modeling and binding site prediction of the NLPA protein. (A) AlphaFold-generated three-dimensional structure of NLPA. (B) Ramachandran plot showing the distribution of residues in favored and allowed regions. (C) Site Map analysis highlighting the predicted druggable binding pocket.

### b. Assessment of druggability across predicted binding sites

Since empirically defined structure or active-site information for NLPA are not available, we relied on the Site Map module of the Schrodinger software. The site-map tool generated five potential binding sites for the NLPA protein (Table 1). Out of the 5 sites generated, the site with a site-score value of 1.109, a D-score value of 0.865, a binding site volume of 127.253, a hydrophilic score of 1.724, and a hydrophobic score of 0.169 was chosen as the best site, i.e., site 1 (Figure 1c). The receptor grid was generated by centering the grid box on the predicted binding site by applying the X, Y, and Z coordinates as 3.09, - 5.56, and -2.31 Å, respectively, and retaining the other parameters at their default values.

**Table 1:** Sitemap data analysis – Results overview of NLPA.

| S. No | Site Name | Site score | Size | Dscore | Volume | Residues |
| --- | --- | --- | --- | --- | --- | --- |
| 1 | Site - 1 | 1.109 | 80 | 0.865 | 127.253 | 47,50,78,95,96,97,100,119,121,150,153,194,210,212,213,238,240,278 |
| 2 | Site - 2 | 0.887 | 71 | 0.897 | 199.283 | 76,77,78,79,82,100,103,104,106,107,108,146,147,148,149,174,193,194,195,213 |
| 3 | Site - 3 | 0.729 | 30 | 0.71 | 82.32 | 120,231,232,233,235,236,239,281,283,287,288 |
| 4 | Site - 4 | 0.706 | 38 | 0.66 | 179.389 | 98,101,102,105,113,114,246,249,285,288,289,291,292 |
| 5 | Site - 5 | 0.645 | 31 | 0.595 | 74.088 | 115,116,117,255,258,259,262,282,283,284,285,286 |

### c. Identification of lead molecules using virtual screening pipelines

An extensive compound collection of nearly 1.6 million small molecules was curated from various chemical libraries for subsequent virtual screening analyses for identifying inhibitors against *A. baumannii* NLPA. The Lig Prep module processed each ligand to produce a single optimized conformer, which was then taken to Glide for virtual screening. The virtual screening workflow (VSW) featured a phased filtering workflow that included HTVS, SP, and XP docking. Final binding poses created during the XP stage were evaluated and ranked according to their Glide XP Gscores. Table 2 lists the top 10 candidates found using hierarchical screening, with binding energies ranging from − 9.155 to − 6.299 kcal/mol.

**Table 2:**
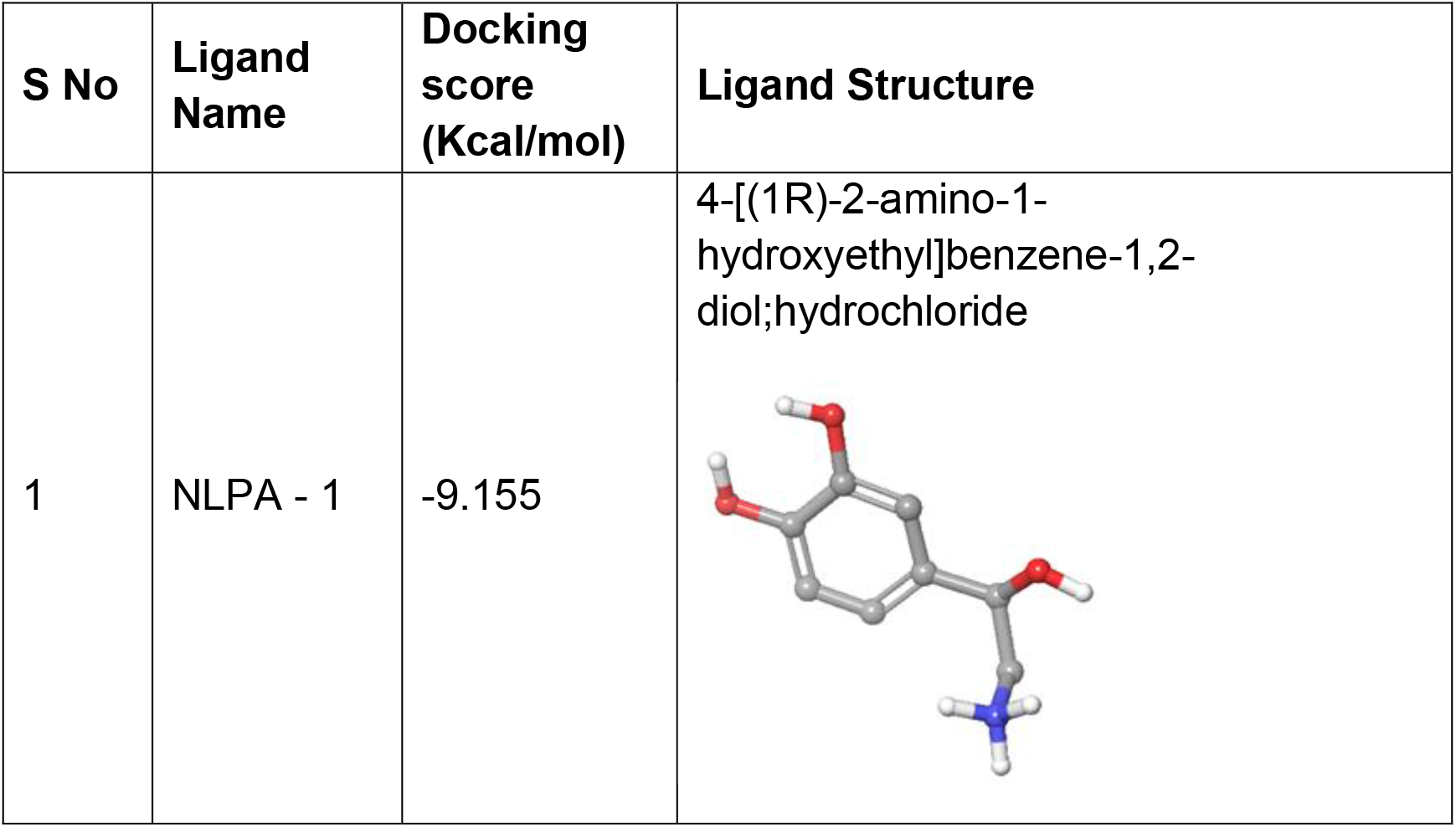
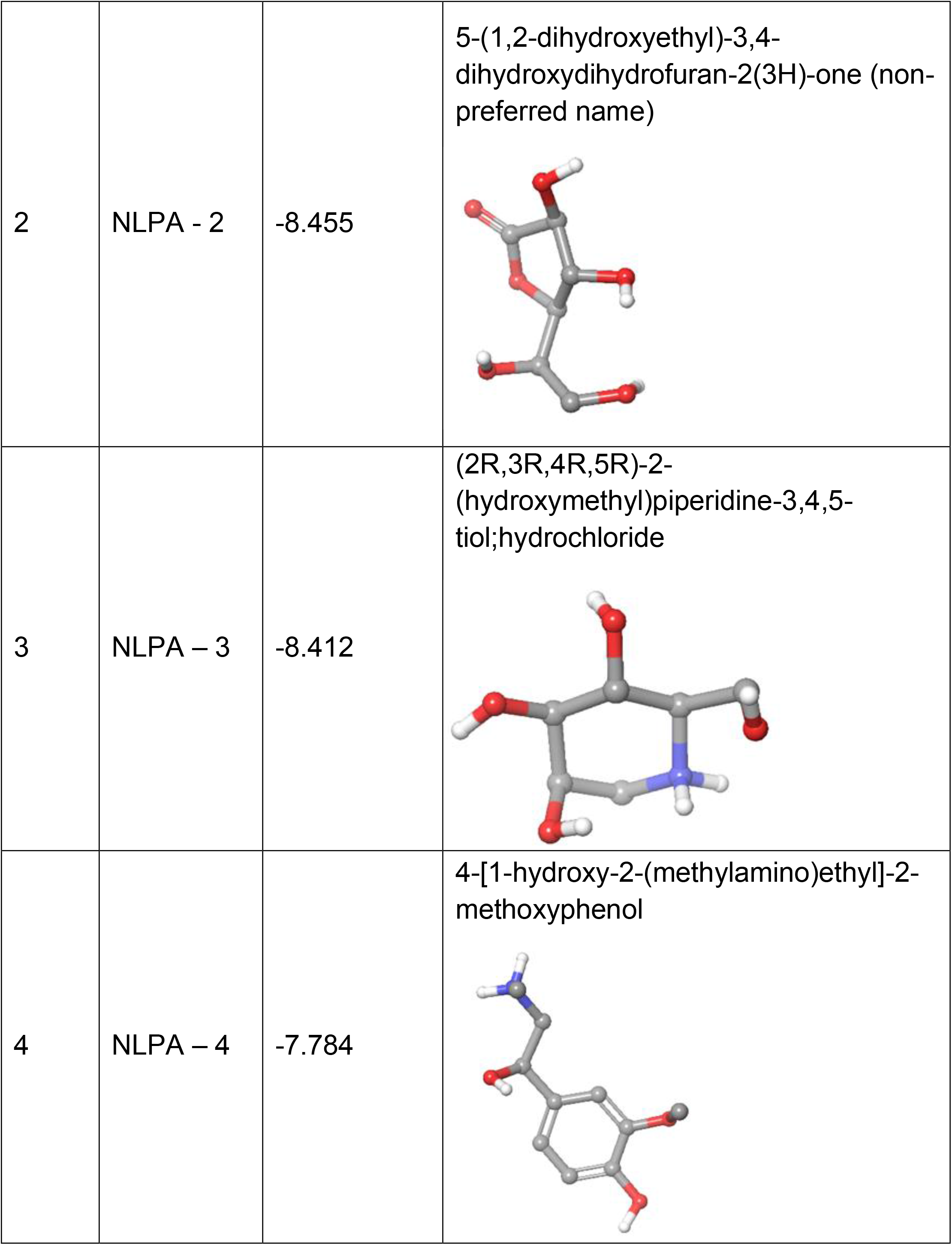
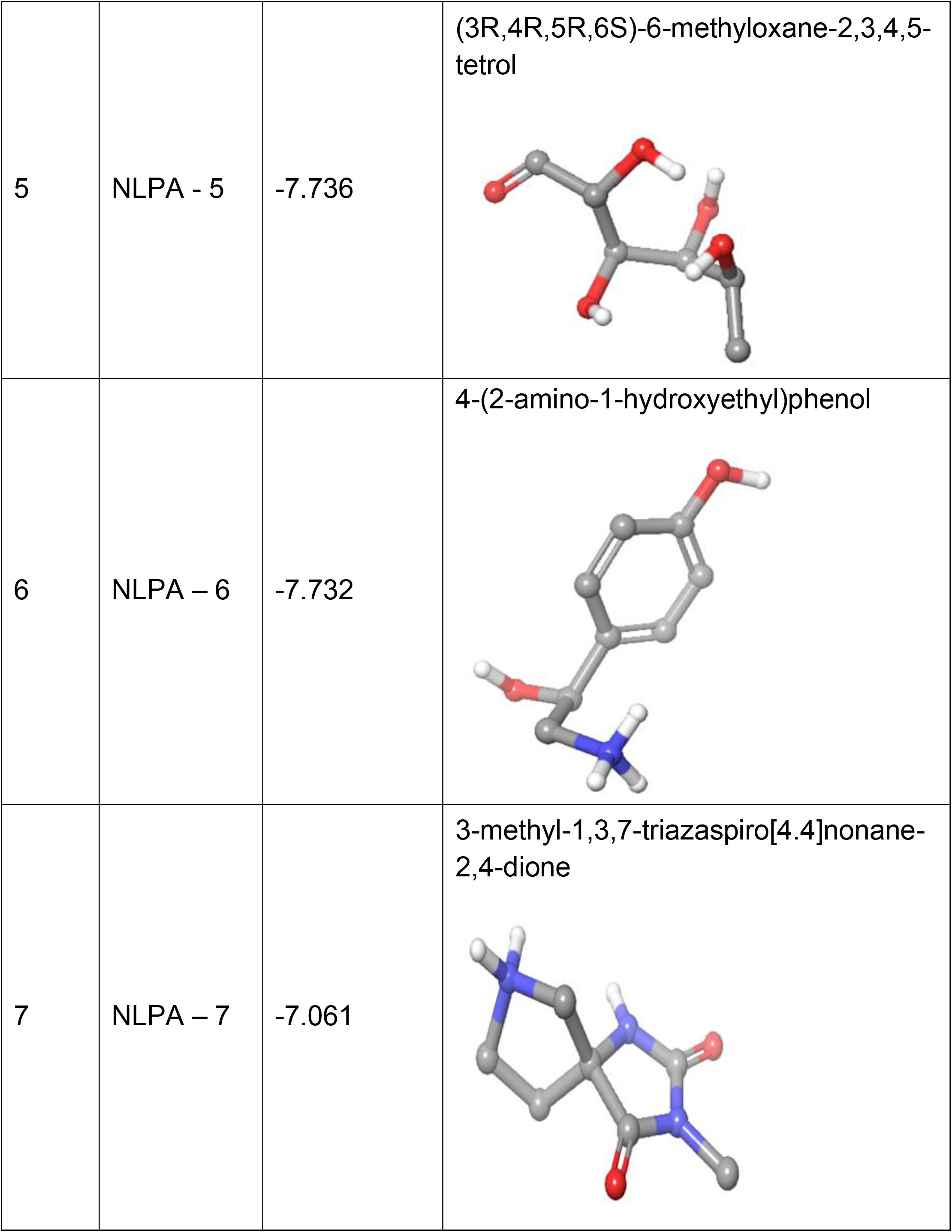
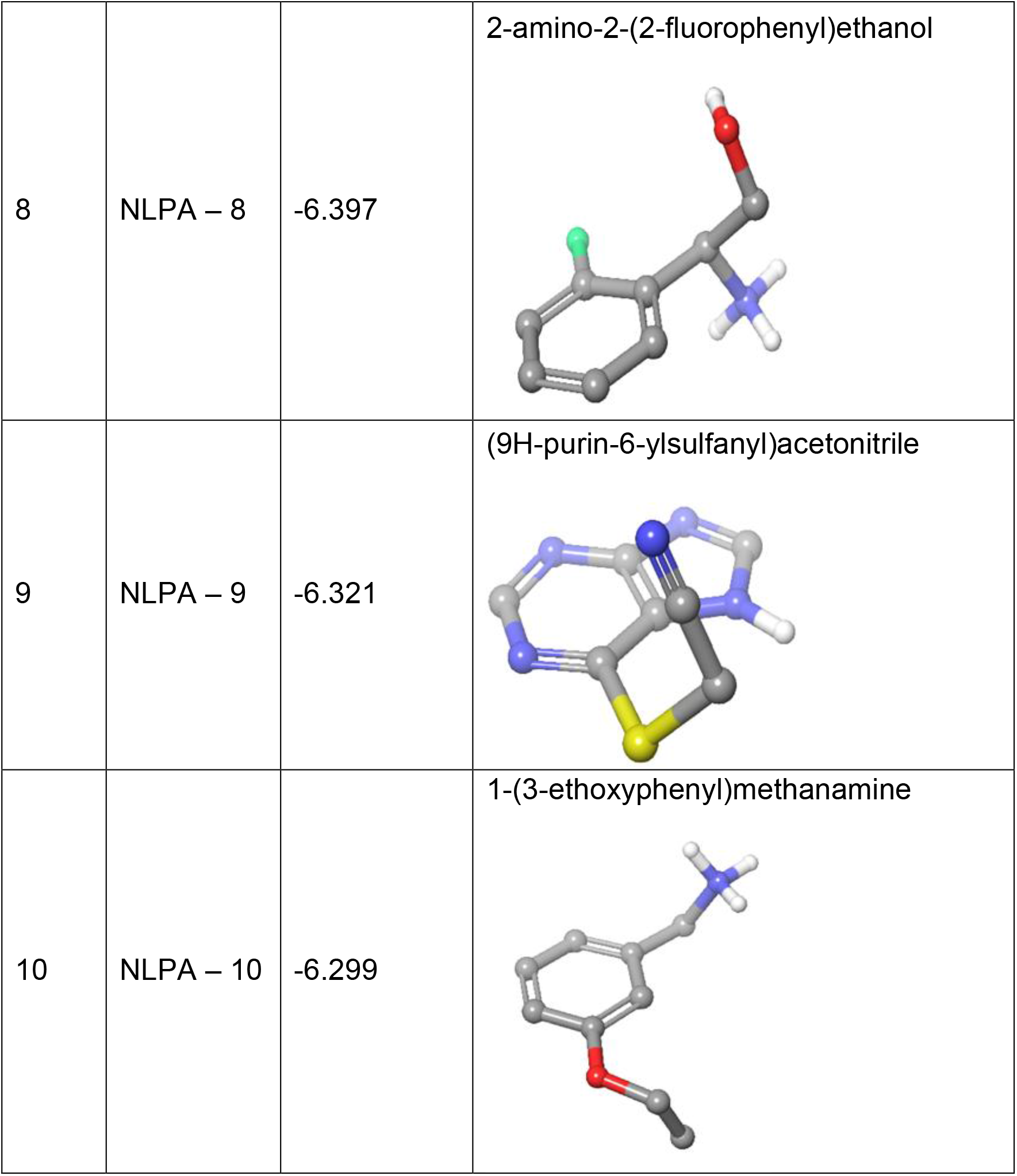
Docking scores of top hits of NLPA.

| S No | Ligand Name | Docking score (Kcal/mol) | Ligand Structure |
| --- | --- | --- | --- |
| 1 | NLPA - 1 | -9.155 | 4-[(1R)-2-amino-1-hydroxyethyl]benzene-1,2-diol;hydrochloride |
| 2 | NLPA - 2 | -8.455 | 5-(1,2-dihydroxyethyl)-3,4-dihydroxydihydrofuran-2(3H)-one (non-preferred name) |
| 3 | NLPA – 3 | -8.412 | (2R,3R,4R,5R)-2-(hydroxymethyl)piperidine-3,4,5-tiol;hydrochloride |
| 4 | NLPA – 4 | -7.784 | 4-[1-hydroxy-2-(methylamino)ethyl]-2-methoxyphenol |
| 5 | NLPA - 5 | -7.736 | (3R,4R,5R,6S)-6-methyloxane-2,3,4,5-tetrol |
| 6 | NLPA – 6 | -7.732 | 4-(2-amino-1-hydroxyethyl)phenol |
| 7 | NLPA – 7 | -7.061 | 3-methyl-1,3,7-triazaspiro[4.4]nonane-2,4-dione |
| 8 | NLPA – 8 | -6.397 | 2-amino-2-(2-fluorophenyl)ethanol |
| 9 | NLPA – 9 | -6.321 | (9H-purin-6-ylsulfanyl)acetonitrile |
| 10 | NLPA – 10 | -6.299 | 1-(3-ethoxyphenyl)methanamine |

### d. Prediction of ligand binding energetics

One of the most prevalent approaches for calculating the free energy of protein–ligand interaction continues to be MM/GBSA. This approach strengthens standard scoring functions, which are used to determine the lowest energy poses and thereby impact the overall accuracy of docking predictions, by improving docking outputs (Wang, Sun et al. 2019). In addition, it offers a more accurate assessment of ligand binding affinities. In this study, relative binding free energy calculations for the top-ranked compounds were carried out using the MM/GBSA tool implemented with the VSGB 2.1 solvation model (Banerjee, Kemmler et al. 2024).

The five compounds with the best binding free energy were selected for further interaction studies based on the calculated ΔG_bind values (Table 3). The molecular interactions between NLPA and the selected top five ligands was outlined using the Ligand Interaction Diagram module (Figure 2). Glu121 consistently contributed to ligand stabilization across all top-ranking ligands with the most favourable ΔG_bind values. In particular, Glu121 formed hydrogen bond interactions with NLPA-2, NLPA-3, and NLPA-4, while with NLPA-6 and NLPA-10 exhibited salt-bridge interactions. The calculated free energy values suggest that the lead compounds can form stable and favorable interactions with the amino acid residues lining the active site.

**Table 3:** MM/GBSA profiles for calculating free energy of the top hits of NLPA.

| S<br>N<br>o | Ligan<br>d<br>Name | MM<br>GBSA<br>dG<br>Bind | MM<br>GBSA<br>dG<br>Bind<br>Coulom<br>b | MM<br>GBSA<br>dG<br>Bind<br>Covale<br>nt | MM<br>GBSA<br>dG<br>Bind<br>Hbond | MM<br>GBSA<br>dG<br>Bind<br>Lipo | MM<br>GBSA<br>dG<br>Bind<br>Solv<br>GB | MM<br>GBSA<br>dG<br>Bind<br>vdW |
| --- | --- | --- | --- | --- | --- | --- | --- | --- |
| 1 | NLPA<br>- 6 | -40.26 | -50.94 | 1.53 | -3.05 | -9.28 | 39.35 | -16.04 |
| 2 | NLPA<br>- 3 | -36.7 | -47.57 | 1.62 | -4.01 | -6.79 | 34.83 | -14.78 |
| 3 | NLPA<br>- 4 | -36.55 | -52.11 | 4.07 | -3.32 | -13.88 | 41.57 | -11.72 |
| 4 | NLPA - 10 | -35.97 | -26.57 | 0.17 | -2.08 | -13.75 | 35.59 | -28.33 |
| 5 | NLPA - 2 | -35.38 | -46.54 | 6.91 | -4.79 | -5.27 | 27.4 | -13.09 |

**Figure 2.**
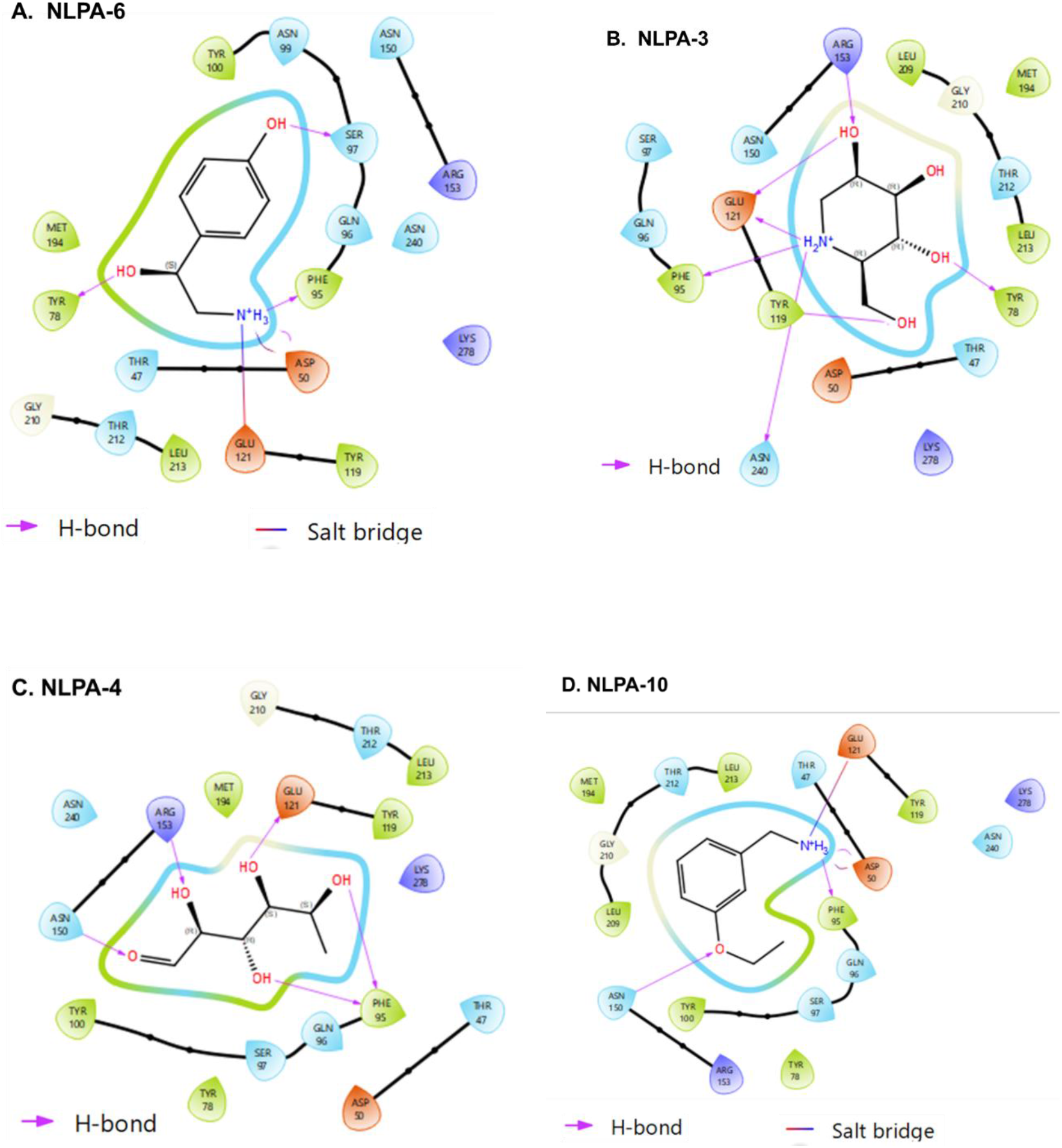
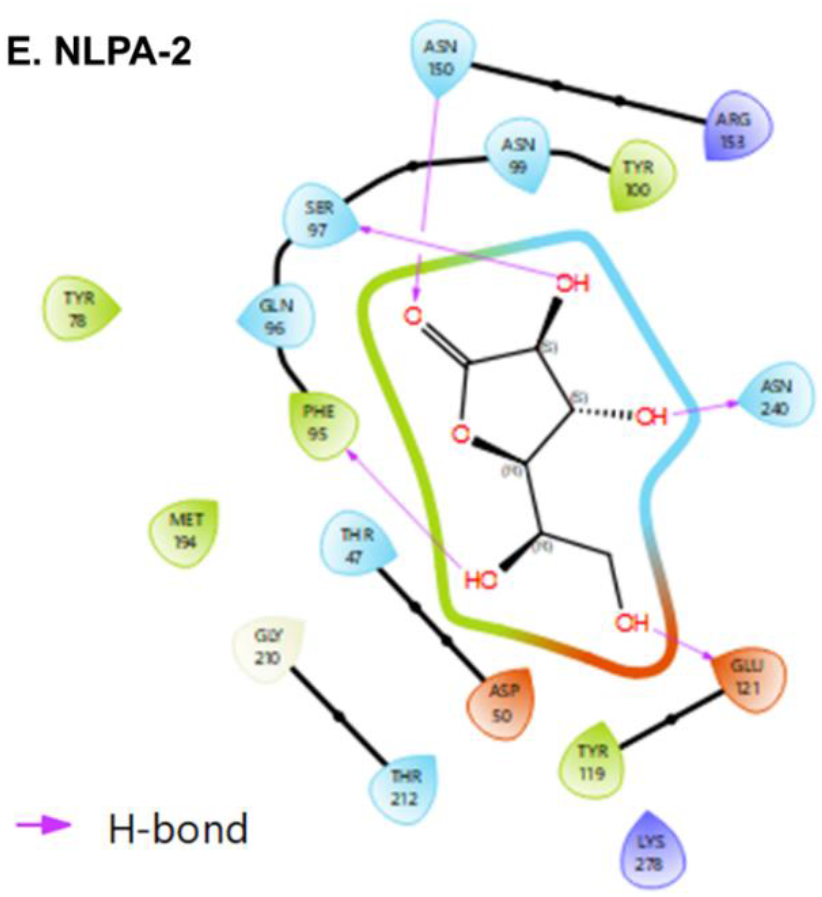
A-E: Ligand interaction diagrams depicting the binding interactions: Molecular interaction diagrams of the top five compounds with key amino acid residues in the predicted NLPA binding pocket.

**Figure 3.**
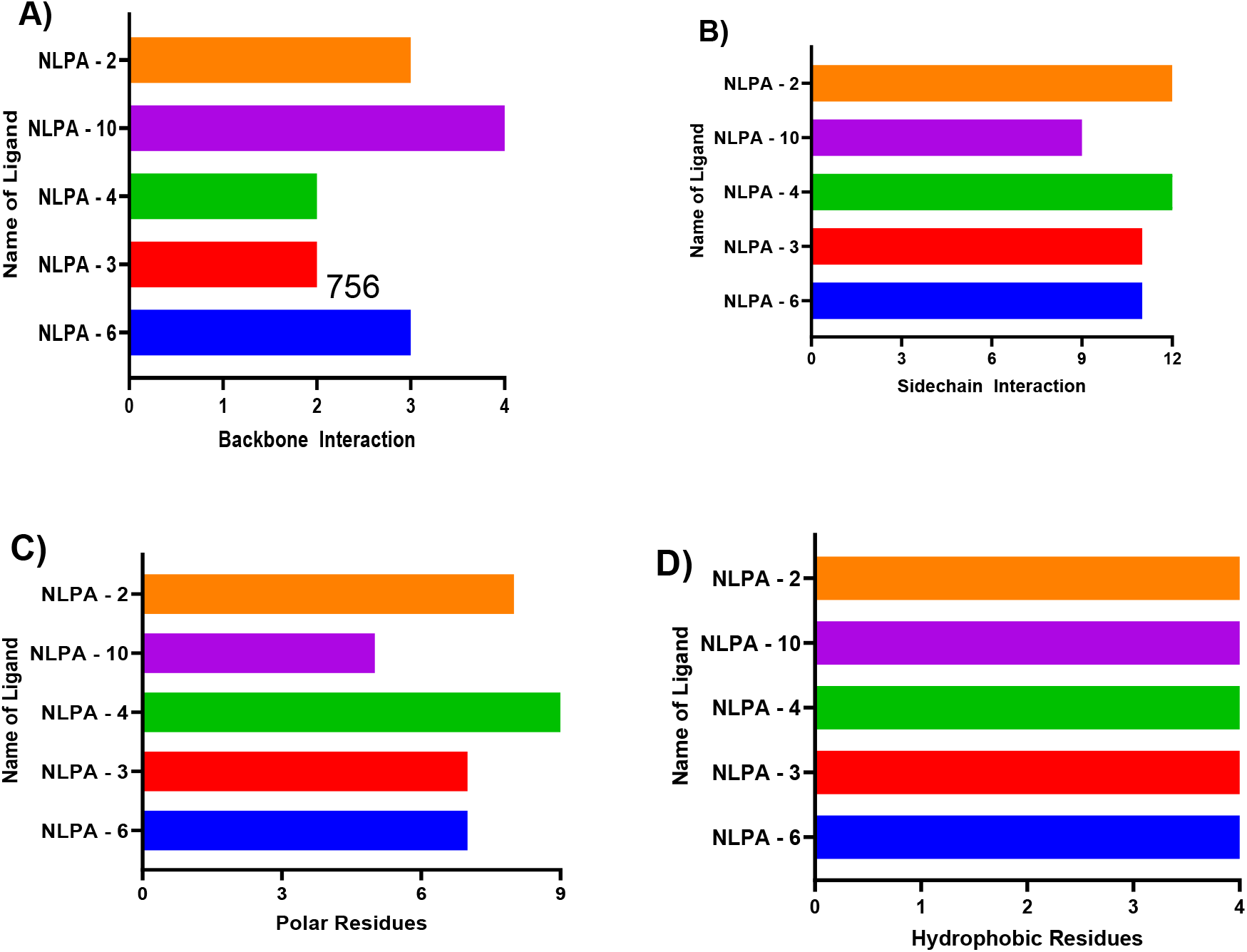
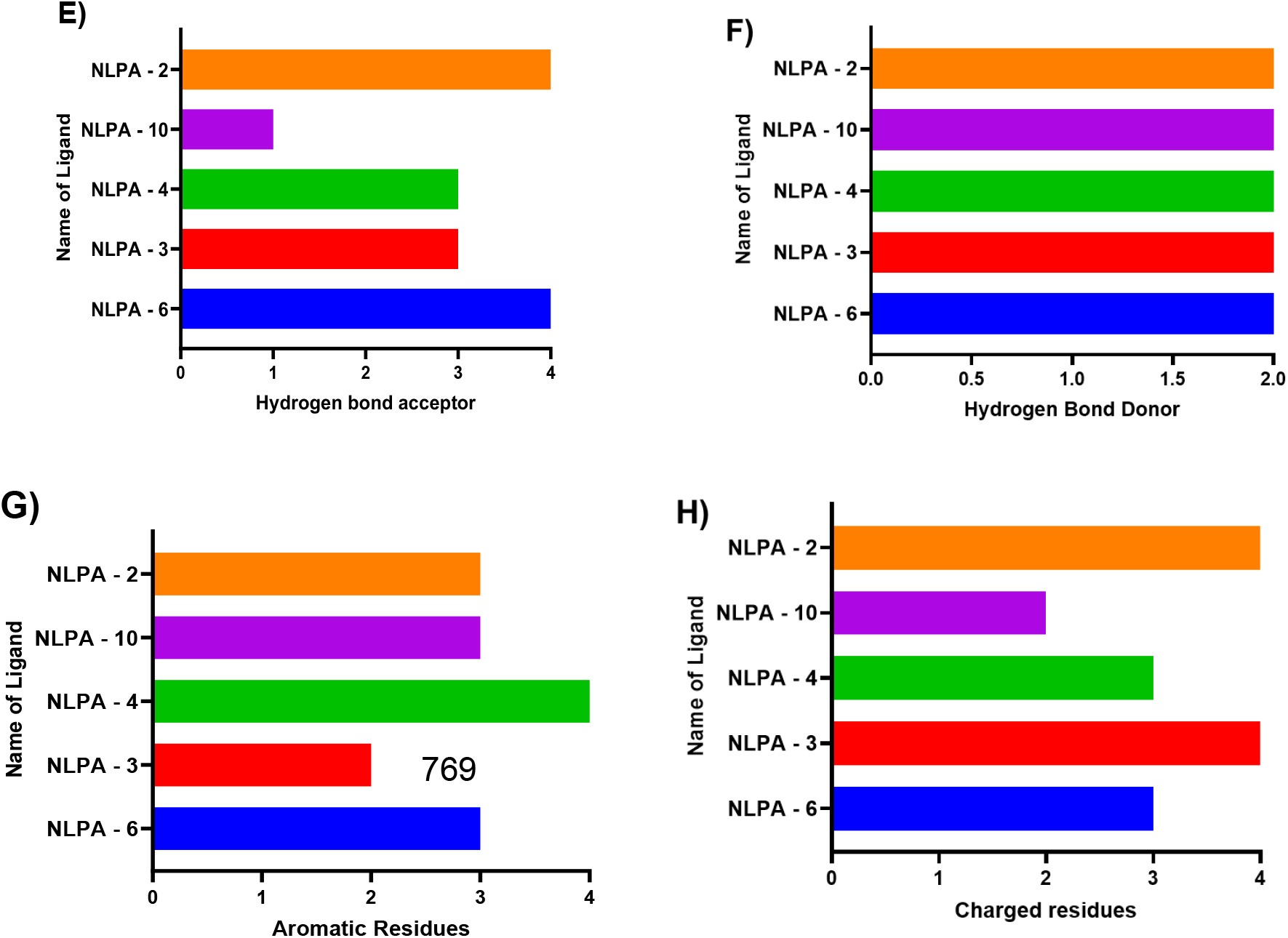
A-H: Interaction fingerprint analysis: Structural fingerprint analysis illustrating the distribution of backbone, side-chain, polar, and hydrophobic residue interactions between the NLPA protein and the top five docked ligands.

The ligand-protein complexes were further stabilized by hydrogen bonding interactions involving a number of nearby residues in addition to Glu121. Arg153 formed hydrogen bonds with NLPA-3 and NLPA-4, while Phe95 interacted with NLPA-2, NLPA-3, NLPA-4, and NLPA-10. Likewise, Tyr78 connected with NLPA-3 and NLPA-6, whereas Asn240 formed hydrogen bonds with NLPA-2 and NLPA-3. Tyr119 and NLPA-3, Ser97 with NLPA-2 and NLPA-6, and Asn150 with NLPA-2, NLPA-4, and NLPA-10 were shown to have additional stabilizing interactions. Together, these interactions imply that a network of hydrogen bonds and electrostatic contacts accommodates the selected ligands inside the binding cavity of NLPA, which probably contributes to the favourable binding energies as shown in the docking and MM-GBSA studies.

### e. Prediction of pharmacokinetic and toxicity parameters

The Qik Prop module was used to assess the ADME parameters of the chosen hit compounds; Table 4 summarizes the findings. Every chemical completely complied with Lipinski’s rule of five. The predicted parameters were assessed against standard cutoffs, including molecular weight (150–650 Da), octanol/water partition coefficient (QPlogPo/w: –2 to 6.5), aqueous solubility (QPlogS: –6.5 to 0.5), Caco-2 permeability (QPPCaco: >25 considered acceptable), MDCK permeability (QPPMDCK: <25 poor, >500 high), human oral absorption (>80% high; <25% poor), and human serum albumin binding (QPlogKhsa: –1 to 1.5).

**Table 4:** Physicochemical and pharmacokinetic profiles of top hit 5 compounds of NLPA.

| S No | Ligand Name | mol MW | Rule Of Five | Percent Human Oral Absorption | QPP Caco | QPP MDCK | QP logK<sub>h</sub><sub>sa</sub> | QP logP<sub>o/w</sub> | QP logS |
| --- | --- | --- | --- | --- | --- | --- | --- | --- | --- |
| 1 | NLPA - 6 | 153.18 | 0 | 56.418 | 79.3 | 35.36 | -0.78 | -0.77 | 0.419 |
| 2 | NLPA - 3 | 163.17 | 0 | 42.619 | 37.8 | 15.86 | -0.84 | -2.14 | 0.236 |
| 3 | NLPA - 4 | 197.23 | 0 | 67.675 | 191 | 91.66 | -0.58 | -0.02 | -0.627 |
| 4 | NLPA - 10 | 151.21 | 0 | 86.724 | 645 | 340.7 | -0.33 | 1.621 | -0.651 |
| 5 | NLPA - 2 | 178.14 | 0 | 44.577 | 52.7 | 20.53 | -0.97 | -2.25 | -0.738 |

All five compounds had desirable drug-like qualities and met Lipinski’s Rule of Five without any violations. The estimated oral absorption in humans was between 42.6% to 86.7%, indicating moderate to high bioavailability. Notably, NLPA-10’s higher QPPCaco and QPPMDCK values showed that it had the greatest expected absorption and permeability. While QPlogPo/w and QPlogS values among the compounds suggested appropriate lipophilicity and water solubility, compounds NLPA-4 and NLPA-6 showed appreciable permeability profiles. All together, these pharmacokinetic factors strengthen the chosen compounds’ being potential for further research.

The ProTox-3 server was used to predict toxicity metrics such as hepatotoxicity, carcinogenicity, mutagenicity, cytotoxicity, and acute oral toxicity (LD_50_) in order to further assess safety. The projected LD_50_ values showed different levels of toxicity among the substances, ranging from 90 to 10700 mg/kg, corresponding to toxicity classes 3–6. While NLPA-3, NLPA-6, and NLPA-10 belonged to toxicity class 4, indicating relatively low toxicity, NLPA-2 had the safest toxicity profile (class 6). The toxicity of NLPA-4 was much greater (class 3) (Table 5). For the most part, the majority of the selected compounds showed acceptable safety profiles according to the projected toxicity endpoints, indicating their potential for additional studies.

**Table 5:** Toxicity profiles of top hit 5 compounds of NLPA.

| S No | Ligand Name | LD<sub>50</sub> (mg/kg) | Toxicity Class | Hepatotoxicity | Carcinogenicity | Cytotoxicity | Mutagenicity |
| --- | --- | --- | --- | --- | --- | --- | --- |
| 1 | NLPA-2 | 10700 | 6 | Inactive | Inactive | Inactive | Inactive |
| 2 | NLPA-3 | 1200 | 4 | Inactive | Inactive | Inactive | Inactive |
| 3 | NLPA-6 | 1090 | 4 | Inactive | Inactive | Inactive | Inactive |
| 4 | NLPA-10 | 313 | 4 | Inactive | Inactive | Inactive | Inactive |
| 5 | NLPA-4 | 90 | 3 | Active | Inactive | Inactive | Inactive |

Based on the integrated assessment of all the above parameters, NLPA-6 illustrated the best binding affinity, while NLPA-3 also showed strong binding interactions which has significant inside the predicted binding pocket. NLPA-3 and NLPA-6 were chosen for additional molecular dynamics simulations to assess the stability and dynamic behaviour of their complexes with NLPA in light of their favourable computational defining features.

### f. Comparative interaction fingerprint assessment

The residue-level interaction patterns between the shortlisted ligands and the NLPA binding pocket were described using interaction fingerprint analysis. Backbone, side-chain, polar, hydrophobic, hydrogen bond acceptor and donor residues, as well as aromatic and charged residues, were among the residue classes with which the analysis measured the number of interactions formed. Interestingly, all of the five compounds showed consistent interactions with a number of important residues in the anticipated binding site. TYR78 and TYR119 contributed via side-chain, hydrophobic, and aromatic contacts, while residues THR50 and THR212 engaged in side-chain, polar, and charged interactions. Furthermore, PHE95 mostly made backbone connections with the ligands, whereas ASN150 and ASN240 participated in side-chain and polar interactions and GLU121 formed polar and charged interactions with the residue. When taken as a whole, these conserved interaction patterns indicate that the shortlisted ligands are stabilized by a mix of polar, aromatic, and hydrophobic interactions and occupy a comparable binding area inside the NLPA predicted binding pocket.

### g. Characterization of system stability during simulations

MD simulations were conducted using the Desmond module in Schrödinger 2022-4 to evaluate the binding stability and dynamic behavior of the protein–ligand complexes at both molecular and atomic scales. Simulations were performed for 100 ns under an isothermal–isobaric (NPT) ensemble to ensure proper equilibration of each complex. System stability was assessed through multiple structural and dynamic parameters, including RMSD, RMSF, hydrogen-bond profiles and radius of gyration (Rg). The RMSD trajectories were analyzed to monitor the complexes’ conformational stability, with consistently low RMSD values indicating that the ligand remained firmly accommodated inside the protein-binding pocket throughout the simulation. This also gives a quantitative measure of the structural variation of the protein backbone during the simulation. (Thirumal Kumar, Lavanya et al. 2017). According, to the RMSD profiles, both complexes reached equilibrium early in the simulation and were mostly stable throughout the trajectory. The NLPA-3 complex had much smoother stability about ∼1.8–2.1 Å, whereas the NLPA-6 complex exhibited modest fluctuations before stabilizing around ∼2.0–2.4 Å, suggesting stable ligand accommodation inside the binding site for both systems (Figure 4: A and B). The obtained RMSD values revealed slight structural variations in the protein backbone, suggesting that ligand binding did not cause major conformational changes in the NLPA structure.

**Figure 4.**
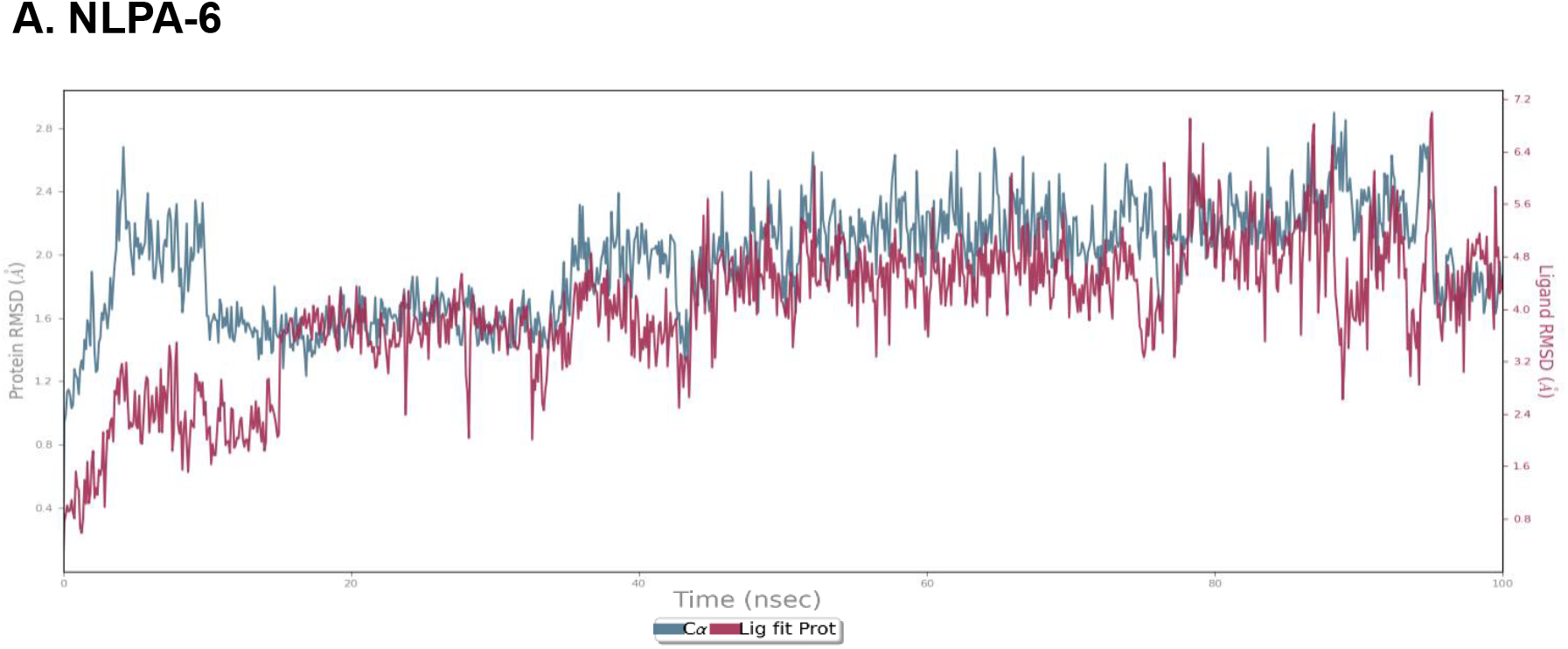
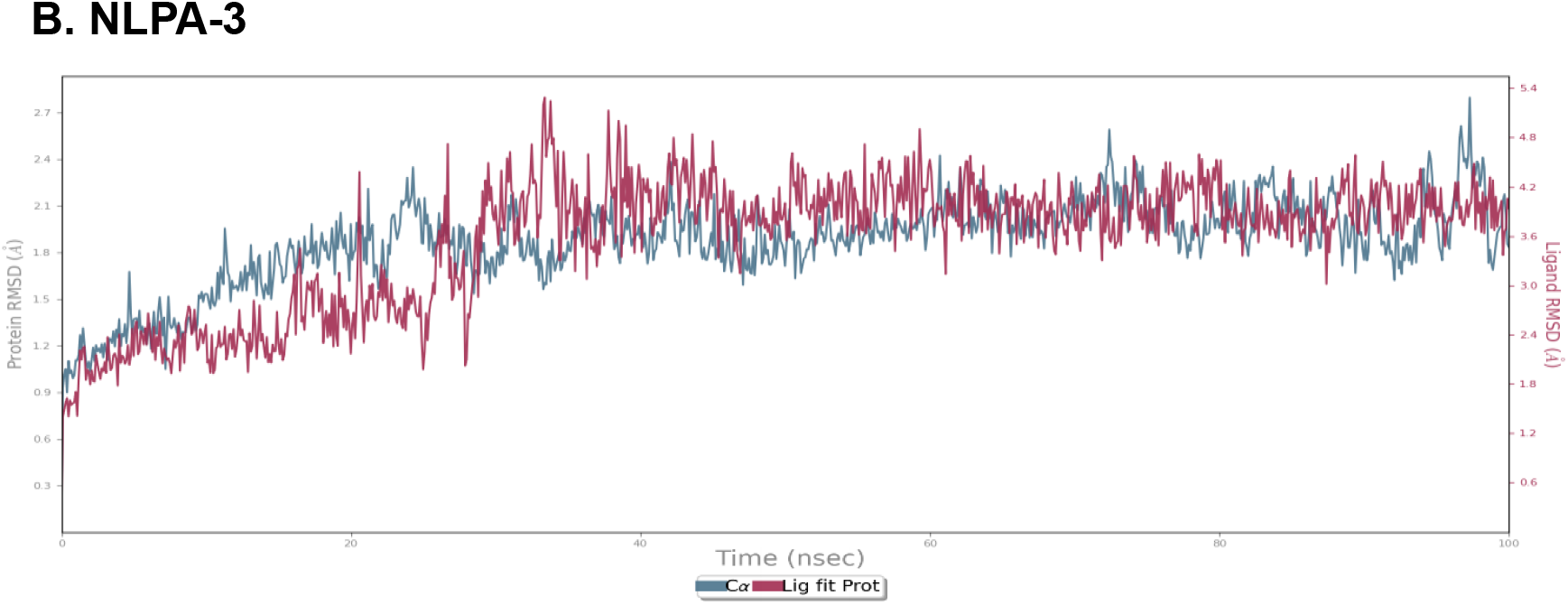
A and B : RMSD profile of the protein backbone and ligand: The protein– ligand complexes over a 100 ns molecular dynamics simulation, NLPA indicating the conformational stability of the complexes.

The RMSF analysis revealed that most residues had variations below ∼1.5 Å, similar to the RMSD findings. Higher flexibility was predominantly found in terminal sections and loop segments, as is characteristic of protein dynamics. Importantly, residues inside the predicted binding pocket had very low RMSF values, indicating that ligand interaction did not alter the protein’s structural integrity (Figure 5: A and B).

**Figure 5.**
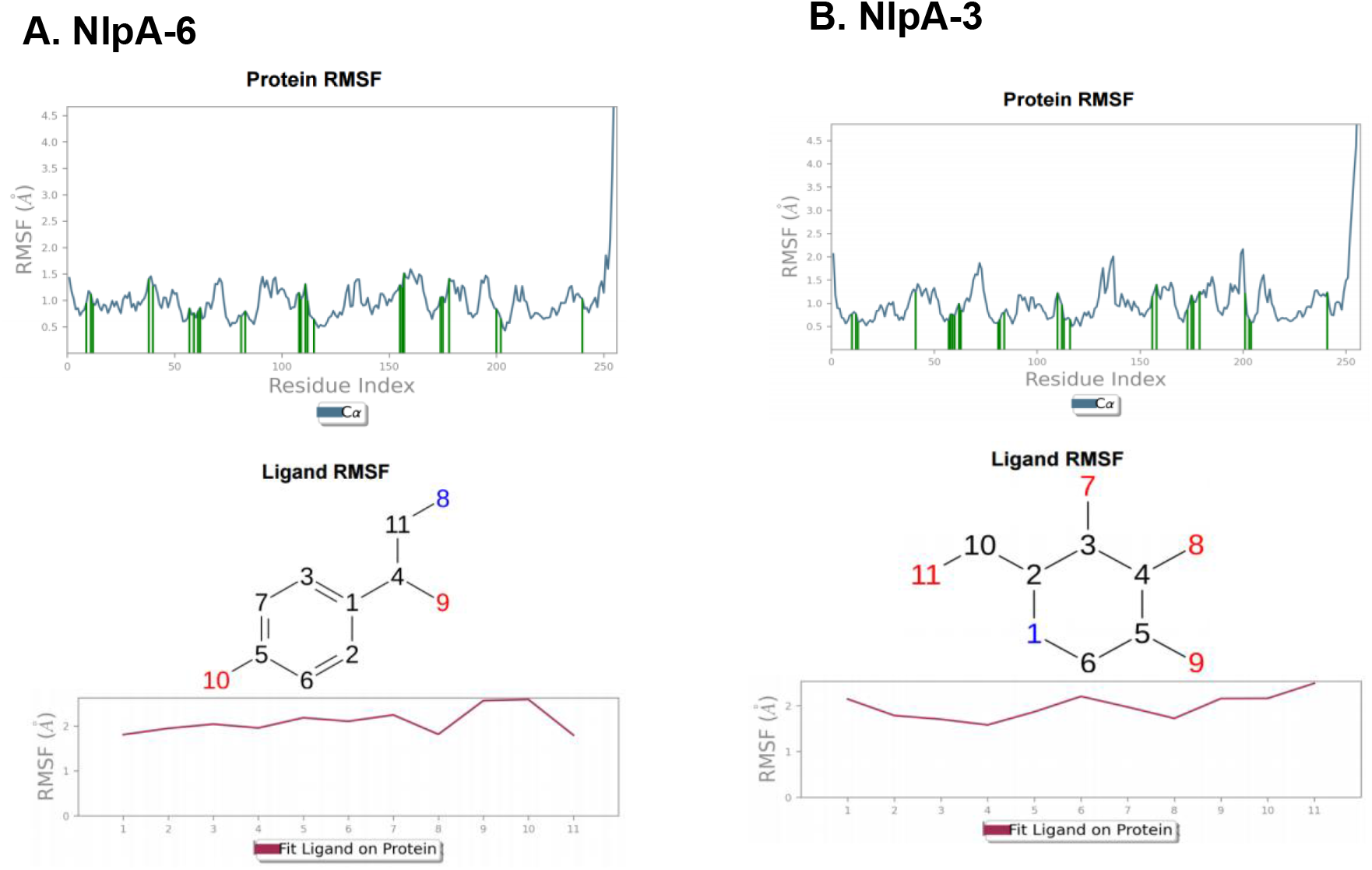
A and B: RMSF analysis: Residue level flexibility analysis of NLPA residues during the 100 ns molecular dynamics simulation, illustrating residue-level flexibility of the protein in the presence of bound ligands.

Hydrogen bonding interactions are essential for stabilizing protein-ligand complexes because they maintain strong and specific contacts inside the binding pocket (Li, Huang et al. 2024). Accordingly, the histogram shows critical interactions between ligands and proteins. In this context, the interaction percentages and amino acid residues indicate the proportion of H-bonds, hydrophobic bonds, ionic interactions, and water bridges observed throughout the simulated trajectory. Consistent with this observation, protein-ligand interactions demonstrated the frequency of contacts between ligands and NLPA residues throughout the simulation. Specifically, residues such as Ser49, Tyr78, Phe95, Glu121, and Asp147 had large interaction percentages with the NLPA-6 complex, mostly via hydrogen bonding and hydrophobic contacts, although occasional water-mediated bridges also helped to stabilize it. Similarly, Ser49, Glu121, Tyr119, and Asn240 were identified as important interacting residues in the NLPA-3 complex, with hydrogen bonding being the major interaction type, reinforced by additional hydrophobic and water-bridge interactions (Figure 6: A and B).

**Figure 6.**
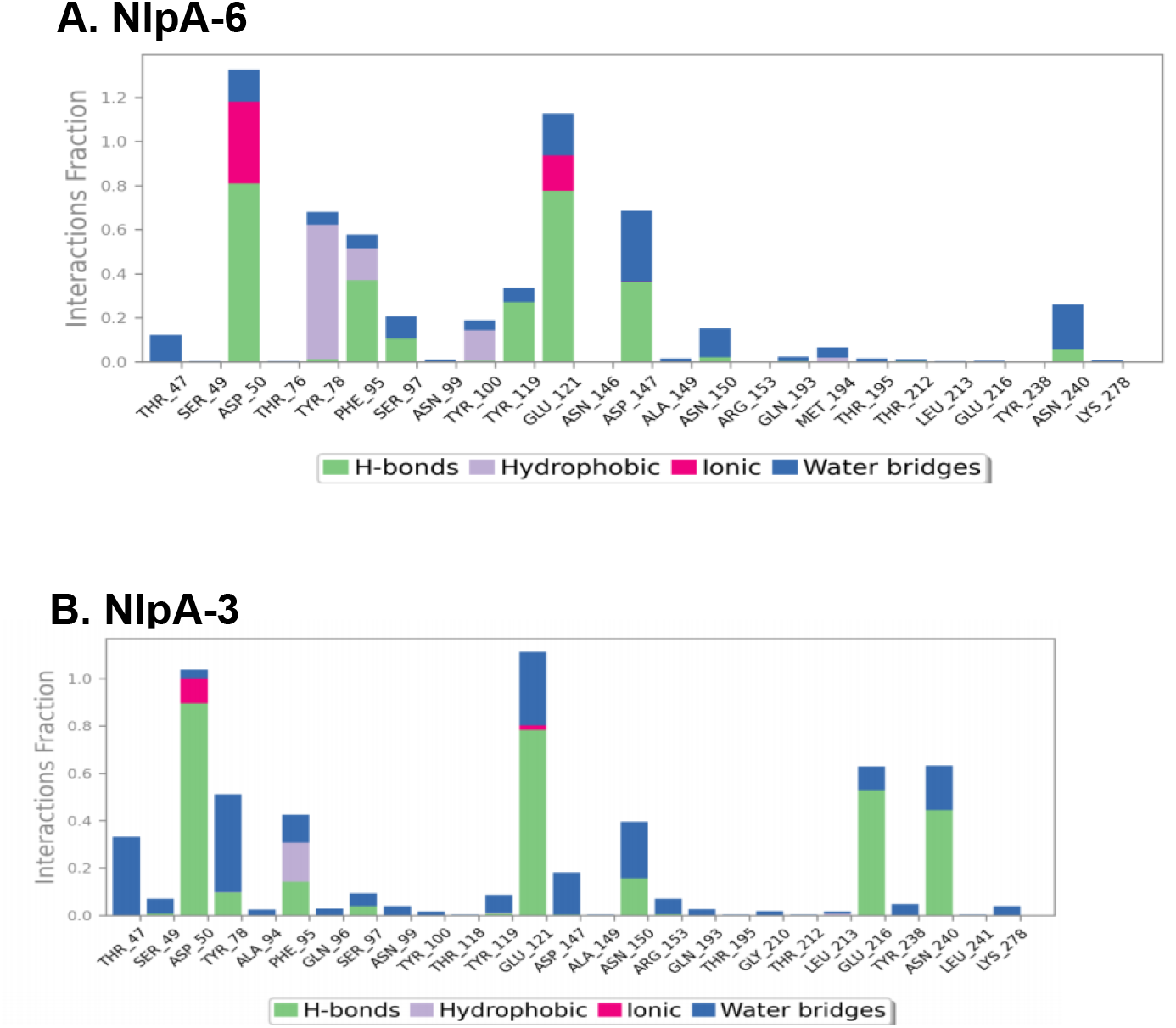
A and B: Histogram analysis of the ligands: Residue interactions showing the fraction of hydrogen bonds, hydrophobic contacts, ionic interactions, and water bridges during the simulation trajectory of NLPA.

The interaction timeline graphs demonstrated the durability of these interactions throughout the simulation period. In the NLPA-6 complex, residues Ser49 and Glu121 maintained constant connections over the majority of the simulation time, whereas Tyr78, Tyr119, and Asn240 added additional transient contacts. Similarly, in the NLPA-3 complex, Ser49, Glu121, and Tyr119 maintained connections throughout the simulation, emphasizing their importance in maintaining the ligand inside the binding cavity (Figure 7: A and B).

**Figure 7.**
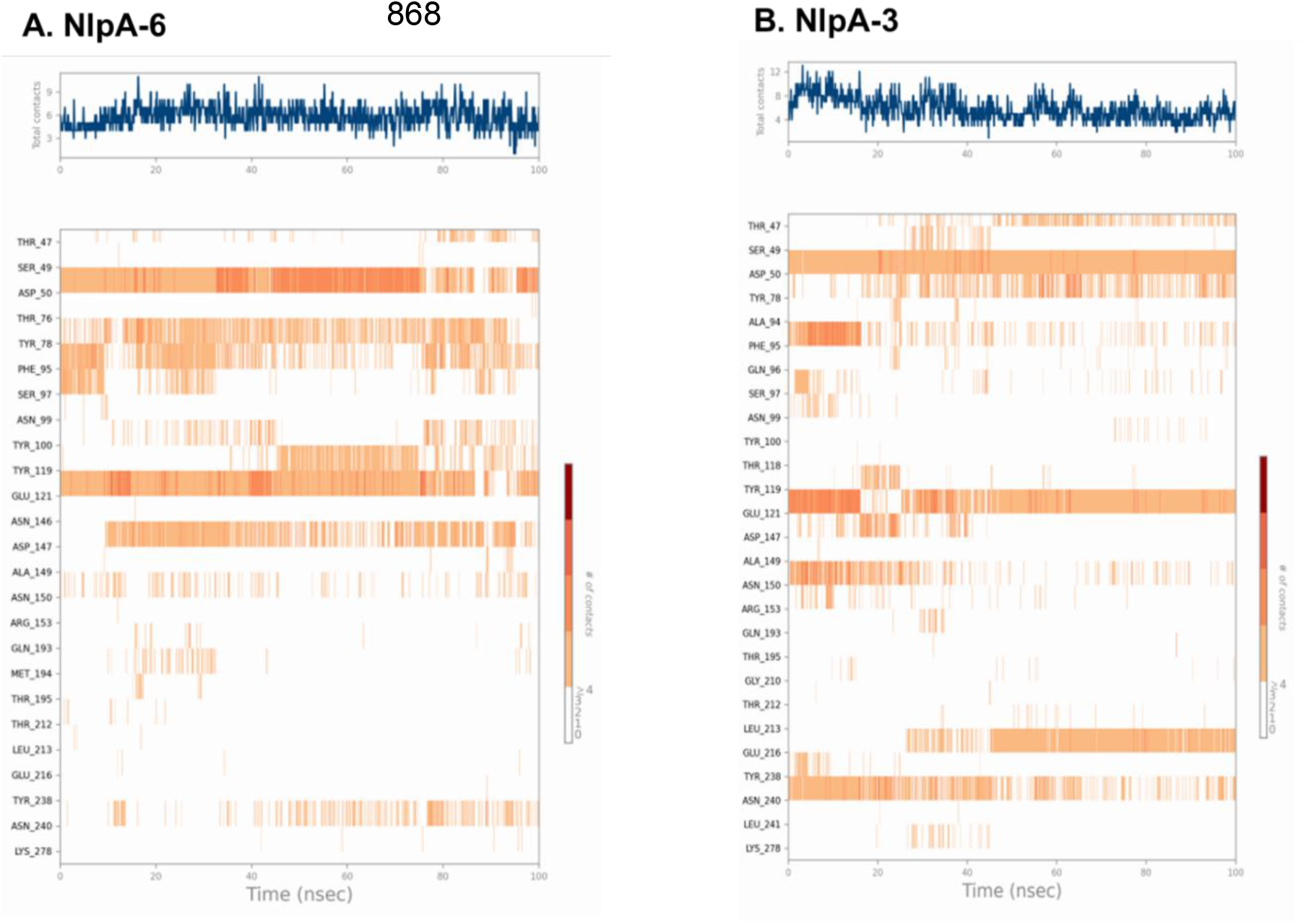
A and B: interaction timeline graphs: Timeline representation of ligand– NLPA residue interactions illustrating the persistence and dynamics of hydrogen bonding, hydrophobic contacts, ionic interactions, and water-mediated bridges throughout the 100 ns molecular dynamics simulation.

Overall, the simulated interaction analysis revealed that both NLPA-6 and NLPA-3 retain stable binding conformations inside the NLPA active site over the 100 ns simulation, which is supported by persistent hydrogen bonding networks and complementary hydrophobic interactions. These results support the favourable binding energies observed from docking and MM-GBSA calculations, demonstrating the ligands’ stable accommodation inside the NLPA binding pocket.

### h. Correlation mapping of residue motions

The dynamic cross-correlation matrix (DCCM) analysis was used to look into the coordinated residue movements of residues within the NLPA protein during 100 ns molecular dynamics simulations of the NLPA-6 and NLPA-3 complexes. Positively correlated motions (blue regions) were observed in the N-terminal region (residues 1-70) and a central segment (residues 140-180) of the NLPA-6 complex. These movements may contribute to the protein-ligand complex’s structural stability. In contrast, sporadic anti-correlated motions (red patches) were found between distant residue segments, indicating compensatory movements that contribute to overall structural equilibrium throughout the simulation. A similar pattern of correlated movements was seen in the NLPA-3 complex, with strong positively correlated clusters near the N-terminal region (1-60) and core domains at ∼110-150 and ∼160-190 residues. These areas showed more cooperative residue movements than the NLPA-6 complex, suggesting improved internal communication within the protein structure after ligand binding (Figure 8: A and B). Overall, both complexes showed well-defined correlated motion networks, suggesting that ligand binding does not destabilize the protein but rather promotes coordinated residue dynamics that enable sustained protein-ligand interactions over the simulation trajectory.

**Figure 8.**
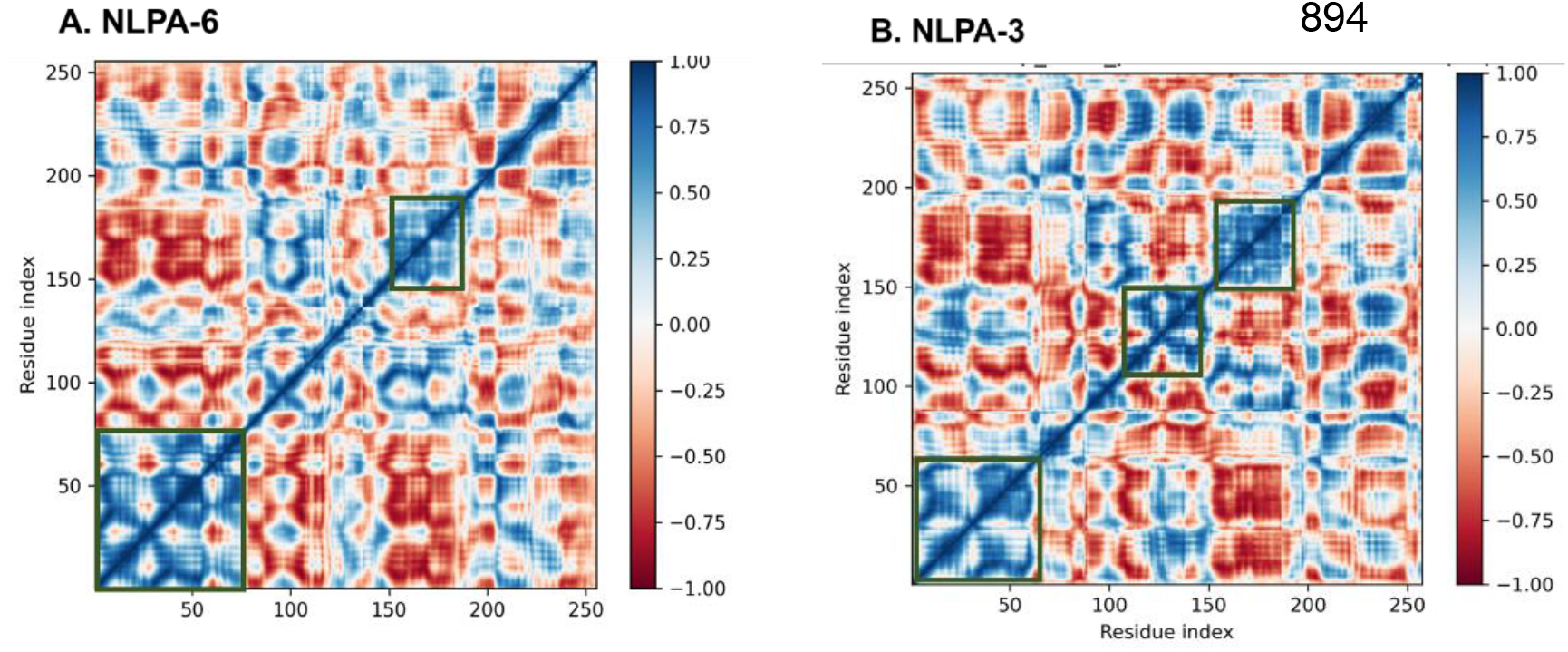
A and B: Correlation mapping: Dynamic cross-correlation matrix plots representing residue–residue motion correlations in NLPA during the 100 ns molecular dynamics simulation for NLPA–6 and NLPA-3 complexes.

**Figure 9.**
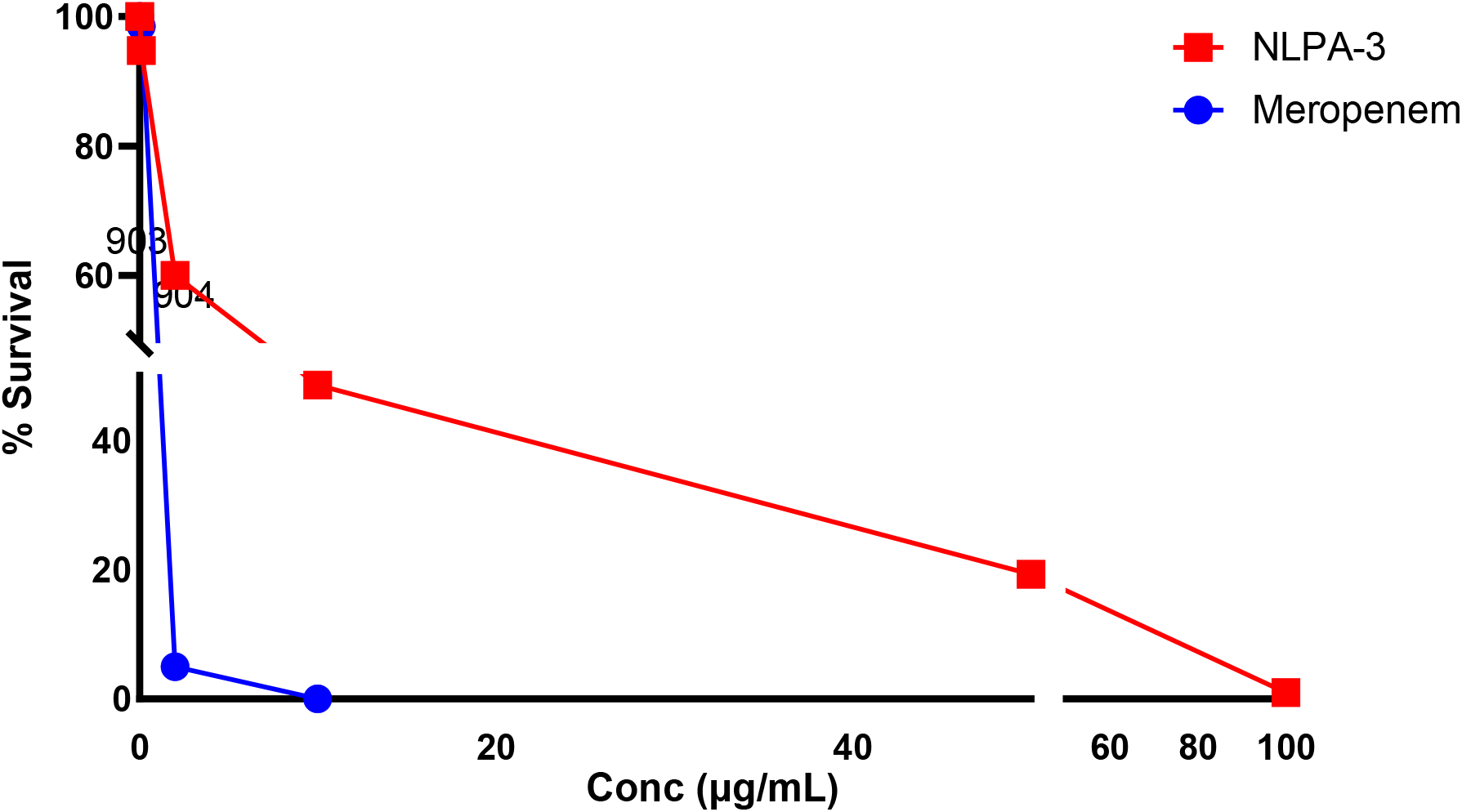
Intracellular antibacterial activity of NLPA-3 against *A*.*baumannii*. Intracellular bacterial survival was evaluated following treatment with varying concentrations of NLPA-3 and compared with meropenem. Results are expressed as percentage bacterial survival relative to untreated control cells.

### i. Antimicrobial and biofilm studies of NLPA-3

NLPA-3 was procured and evaluated in vitro for experimental validation, however NLPA-6 was not accessible from commercial sources. NLPA-3 was subjected to antimicrobial susceptibility and biofilm inhibition against *A. baumannii*. The compound had a minimum inhibitory concentration (MIC) of 125 µg/mL.

The antibiofilm activity of NLPA-3 against *A. baumannii* BAA-747 was evaluated by measuring the percentage inhibition of biofilm formation at different concentrations (1x, 2x and 4x). It inhibited biofilm formation concentration-dependently, starting at 29% at 1x MIC and rising to 53.25% and 55.75% at 2x and 4x MIC, respectively. This dose-responsive pattern suggests biofilm growth is being interfered by this compound. Although the antibacterial efficacy was modest, the observed antibiofilm impact is consistent with anti-virulence techniques that seek to reduce pathogenic characteristics rather than impose direct bactericidal pressure. Biofilm-associated resilience and persistence mechanisms provide relevance of the effectiveness of such methods in *A. baumannii* infections (Rasko and Sperandio 2010). The ability of NLPA-3 to inhibit biofilm formation may be linked to impairment of NLPA-mediated stress adaption and biofilm maintenance pathways. These results, together with the reported persistent binding interactions in silico, suggest NLPA-3’s potential as a candidate for future exploration.

### j. Intracellular activity of NLPA-3 in *A*.*baumannii*

In RAW 264.7 macrophages, NLPA-3’s intracellular antibacterial activity against *A. baumannii* BAA-747 was assessed and compared with standard antibiotic (meropenem). With intracellular survival reducing to 48.5% at 10 µg/mL and 19.25% at 50 µg/mL, NLPA-3 showed concentration-dependent intracellular bacterial decrease. Significant intracellular bacterial inhibition was notably seen at doses below the MIC value (125 µg/mL), suggesting NLPA-3’s capacity to reduce *A. baumannii* intracellular persistence. In comparison, meropenem exhibited complete intracellular bacterial clearance at concentrations ≥10 µg/mL. The cytotoxicity of NLPA-3 was assessed for 24 hours at concentrations ranging from 0 to 1250 µg/mL. The survival percentage of RAW 264.7 cells was 73.53% at 312.5 µg/mL (3x MIC) and dropped to 54.07% at 1250 µg/mL (10x MIC), suggesting that NLPA-3 has moderate concentration-dependent cytotoxicity.

Persistence, immune evasion, and chronic infection are all greatly influenced *by A. baumannii’s* capacity to live within macrophages (Rasko and Sperandio 2010). The observed reduction in intracellular bacterial survival by NLPA-3 indicates its potential to interfere with virulence-associated pathways involved in stress adaptation and host-associated persistence. While NLPA-3’s intracellular antibacterial activity was less than that of meropenem, its concentration-dependent intracellular clearance is consistent with anti-virulence approaches that target bacterial survival and persistence rather than immediate bactericidal effects (Harding, Hennon et al. 2018). Targeting NLPA-associated pathways in *A. baumannii* is therapeutically relevant, as shown by the intracellular activity seen in this work, as well as the antibiofilm effects and stable protein–ligand interactions found by molecular dynamics simulations.

## 4. Conclusion

The NLPA lipoprotein of *A. baumannii* has been studied as prospective anti-virulence target in the current study using an integrated computational method. NLPA-6 and NlLPA-3 were shown to be stable binders with acceptable pharmacokinetic and toxicological profiles by virtual screening, molecular docking, MM-GBSA, and molecular dynamics simulations. NLPA-3’s in vitro validation showed moderate antibacterial activity in addition to concentration-dependent antibiofilm and intracellular inhibitory effects, at sub-MIC concentrations, indicating interference with virulence pathways linked to persistence. Taken together, our results demonstrate the therapeutic value of focusing on NLPA to counteract biofilm-associated persistence in multidrug-resistant *A. baumannii*. Future studies focusing on structural optimization and analog development of NLPA-3 could aid in further improvement of its potency and therapeutic efficacy against *A. baumannii*.

## Acknowledgements

We acknowledge the support of the National Institute of Pharmaceutical Education and Research, Hyderabad, and the Department of Pharmaceuticals (DoP), Ministry of Chemicals and Fertilizer. A part of the study was supported by ICMR small grant IIRP 4907. UB thanks ICMR-SRF grant (AMR/Fellowship/23/2022-ECD-ll).

## Ethical approval

Not required

## Data availability

Not applicable

